# Spatio-Temporal Recruitment of Adult Neural Stem Cells for Transient Neurogenesis During Pregnancy

**DOI:** 10.1101/2021.07.11.451957

**Authors:** Zayna Chaker, Corina Segalada, Fiona Doetsch

**Affiliations:** Biozentrum, University of Basel, Basel, Switzerland

**Author notes:** **Corresponding author:** Fiona Doetsch, Biozentrum, University of Basel, Klingelbergstrasse 50/70, CH 4056 Basel, Switzerland. These authors contributed equally to this work (Zayna Chaker, Corina Segalada).

## Abstract

Neural stem cells (NSCs) in the adult mouse brain contribute to lifelong brain plasticity. NSCs in the adult ventricular-subventricular zone (V-SVZ) are heterogeneous and, depending on their location in the niche, give rise to different subtypes of olfactory bulb interneurons. Here, we show that during pregnancy multiple regionally-distinct NSCs are dynamically recruited at different times. Coordinated temporal activation of these NSC pools generates sequential waves of short-lived olfactory bulb interneuron subtypes that mature in the mother around birth and in the perinatal care period. Concomitant with neuronal addition, oligodendrocyte progenitors also transiently increase in the olfactory bulb. Thus, life experiences, such as pregnancy, can trigger transient neurogenesis and gliogenesis under tight spatial and temporal control, and may provide a novel substrate for brain plasticity in anticipation of temporary physiological demand.

## Main text

Stem cells in the adult mouse brain dynamically integrate and respond to environmental signals lifelong (*1, 2*). V-SVZ NSCs residing along the lateral ventricles are radial glial fibrillary-acidic protein (GFAP) expressing cells and are found in quiescent or activated states (*2*). NSCs in distinct spatial domains of the V-SVZ give rise to different subtypes of olfactory bulb (OB) interneurons (*2*). Some adult NSCs can be modulated by physiological state. NSCs in the anterior ventral V-SVZ respond to fasting/feeding via the activity of long-range hypothalamic innervation (*3*). However, whether other regionally-distinct pools of stem cells are recruited in different contexts, and whether the timing of their recruitment leads to the addition of specific subtypes of new neurons in the olfactory bulb at physiologically relevant periods is unknown.

The process of becoming a mother consists of a succession of physiological phases, including mating, conception, blastocyst implantation, gestation, birth, perinatal care, lactation, and weaning. Pregnancy increases proliferation in the V-SVZ selectively at gestation day 7 and again at postpartum day (Ppd) 7 (*4*) in a prolactin-dependent manner, which leads to increased neurogenesis in the olfactory bulb (*4, 5*). In addition, perturbing adult V-SVZ neurogenesis results in defects in maternal behavior (*5–8*). However, depending on the timing of manipulation during gestation, and the phase of motherhood at which newborn neurons were examined, differing results were obtained (*5–9*). In light of the discovery of regionally distinct pools of adult neural stem cells, we investigated whether different physiological stages in the process of becoming a mother affects the behavior of spatially distinct neural stem cells, and in turn the dynamics of adult neurogenesis in the olfactory bulb during motherhood.

We first quantified stem cell proliferation (GFAP+ Ki67+) at several timepoints of gestation, and Ppd 7.5 (Fig. 1A). Pregnancy did not evenly enhance proliferation of stem cells, but only of those residing in certain domains. Intriguingly, V-SVZ domains that tend to be more quiescent, such as the ventromedial wall (Fig. 1B and D), the roof (Fig. 1C and E), and the dorsomedial corner (DMC, Fig. S1A) were activated during pregnancy. Stem cells residing in the proliferative dorsolateral wedge (DLW, Fig. 1C and E) and the ventrolateral wall (Fig. 1B and D) were also more recruited during pregnancy. In contrast, the dorsolateral wall and intermediate V-SVZ were not more active during pregnancy (Fig. S1B-D). Notably, each pregnancy-related domain displayed distinct temporal dynamics of recruitment (Fig. 1B, C, F). Dividing NSCs in the roof and the ventral V-SVZ increased at Gd 4.5 (the day of implantation) or Gd7.5, respectively (Fig 1B-C; Fig. S1G). In contrast, in the dorsolateral wedge and in the dorsomedial corner, NSC proliferation had more complex dynamics and increased at several gestation days (Fig. 1C; Fig. S1A). All changes were transient, and stem cell proliferation decreased to virgin levels at Gd12.5 and Gd18.5. Stem cell division in mothers at Ppd 7.5 was similar to virgin females in most domains, with the exception of NSCs in the dorsolateral wedge (Fig. 1C). Transit amplifying cells (GFAP-EGFR+) were also affected in a regional manner with higher proliferation in the ventral V-SVZ (Fig. S1E-F and H). Pregnancy therefore leads to the coordinated recruitment of multiple stem cell pools at specific gestation days in different spatial domains in the V-SVZ (Fig. 1F).

**Fig. 1.**
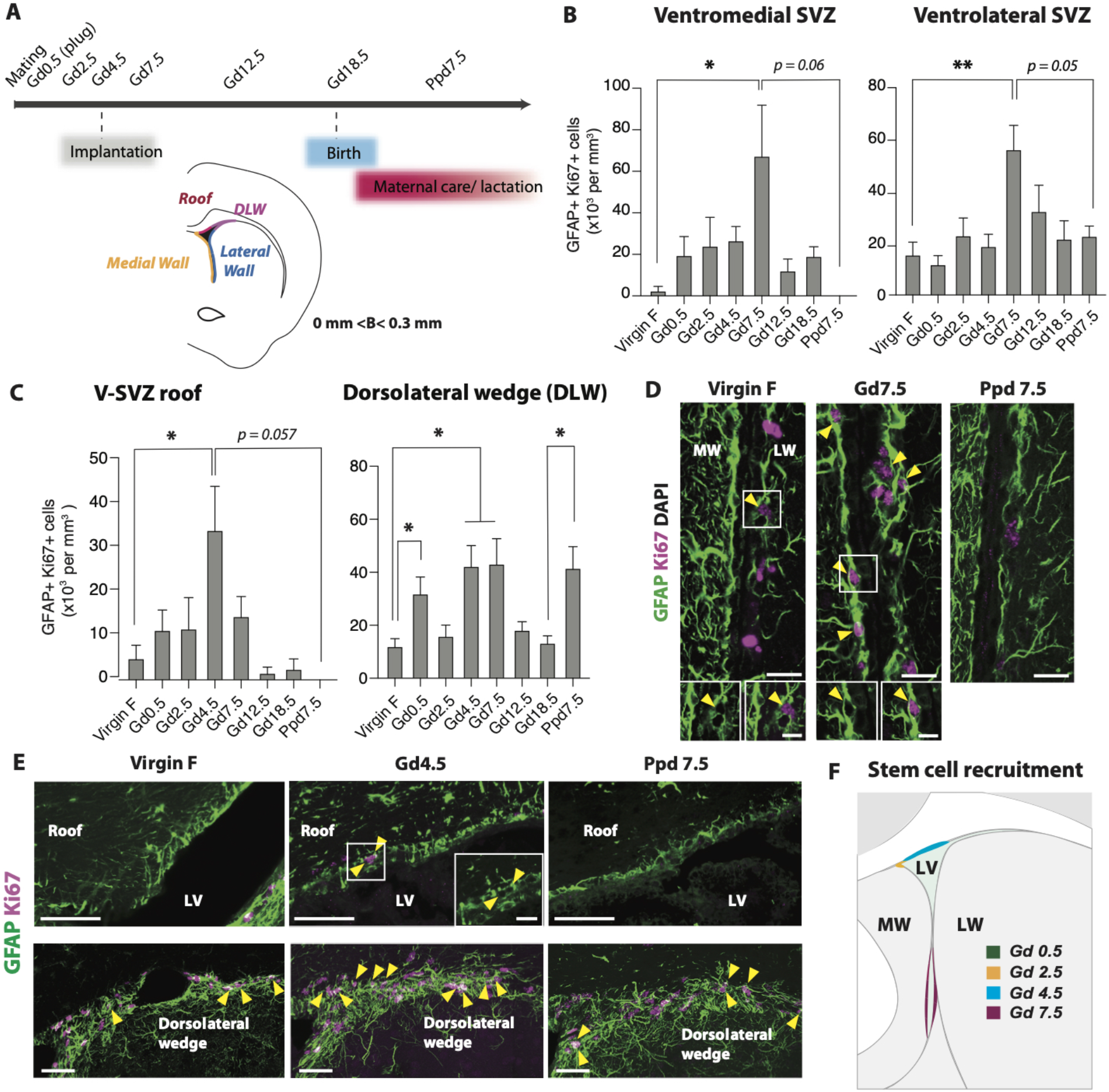
Dynamic spatial and temporal response of V-SVZ NSCs to pregnancy. (**A**) *Top*: Timeline of gestation (Gd) or post-partum (Ppd) days analyzed. *Bottom*: Schema of coronal brain section showing V-SVZ around ventricle (lateral wall and medial wall) and roof and dorsolateral domains analyzed in **B-E**. See also Fig. S1D. (**B** and **C**) Quantification of GFAP+Ki67+ cells in different V-SVZ domains. (**D** and **E**) Representative images of dividing NSCs (arrowheads) (GFAP (green), Ki67 (magenta), DAPI (grey)) in ventrolateral and ventromedial V-SVZ (**D**), roof and dorsolateral wedge (**E**) quantified in **B** and **C**. Scale bars: **D**, 20μm, **E**, 50μm, Boxes in D, E, 10μm. (**F**) Summary schema of temporal and spatial recruitment of V-SVZ stem cell domains during pregnancy. LV: lateral ventricle. MW: medial wall, LW: lateral wall, B, Bregma.

V-SVZ stem cells in different domains generate distinct subtypes of OB interneurons (*1, 2*). Embryonic, postnatal and adult-born interneurons are functionally heterogeneous, with specific subpopulations predominantly generated developmentally or during adulthood (*10–12*). In the adult, most newly-generated neurons integrate into the granule cell layer (GCL) of the main olfactory bulb (MOB) (Fig. 2A), and some differentiate into periglomerular cells (GL layer, Fig. 2A). Rarely, interneurons are added to the mitral cell layer (MCL, Fig. 2A) and external plexiform layer (*13*). Low levels of constitutive adult neurogenesis also occurs in the accessory olfactory bulb (AOB, Fig. 2A) (*14*).

**Fig. 2.**
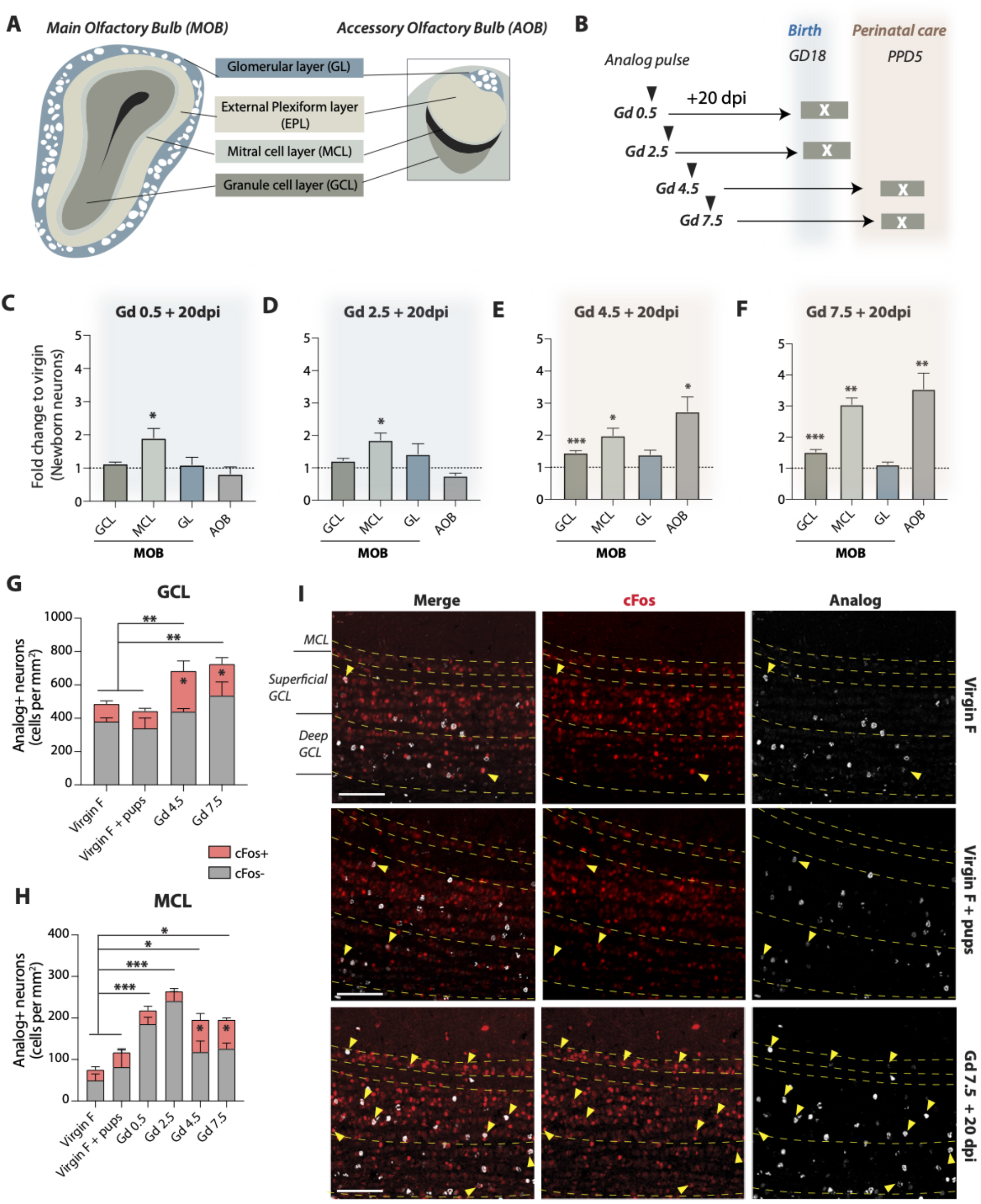
Temporally regulated addition of distinct OB interneuron subtypes born during pregnancy. (**A**) Schema of different layers in MOB and AOB. (**B**) Injection paradigm of pulse time points during pregnancy, and physiological phases after the 20 day-chase period. (**C** to **F**) Quantification of fold-change of newborn neurons (NeuN+ analog+) generated at different gestation days in distinct OB layers compared to matched virgin controls (**G** and **H**). c-fos expression in newborn neurons in the GCL **(G)**, and in the MCL **(H)** of the MOB. Stars inside the pink bar graphs show significant differences in c-fos+ analog+ cell number between virgins and mothers. (**I**) Representative images of analog-labeled cells and c-fos expression in the MOB of mothers pulsed at Gd 7.5, and virgin females with or without pups. Scale bars, 100μm.

To investigate whether the spatial and temporal recruitment of NSCs during pregnancy resulted in the addition of specific interneuron subtypes to the OB at different physiological stages of motherhood, we analyzed olfactory bulbs of mice pulsed once with a thymidine analog on different gestation days. Neurogenesis is typically examined 30 days after labeling, however newly generated neurons are already functionally integrated in the olfactory bulb after two to three weeks (*15*). 20dpi coincides with key physiological events in motherhood, namely parturition for newborn neurons pulsed at Gd0.5 and Gd2.5, and the first week of intense perinatal care and lactation for those generated at Gd4.5 and Gd7.5 (Fig. 2B). We therefore focused first on the 20dpi time point.

The number of newly generated cells within the OB increased in mothers but strikingly their distribution differed depending on their day of birth (Fig. S2A and Fig. S3A). Quantification of thymidine analog and NeuN labeled cells showed that in the GCL in the main olfactory bulb (MOB), newborn neurons only increased in mothers pulsed at Gd4.5 and 7.5 (Fig. 2C to F, Fig. S2B). The increase in Gd 4.5 born interneurons was observed in both the deep and superficial layers, whereas those born at 7.5d were selectively increased in the superficial GCL (Fig. S2C). Moreover, both Calretinin+ and Calretinin-granule cells increased (Fig. S2D and E). This shift in neuronal addition to the superficial GCL is especially interesting, as new neurons are preferentially added to the deep granule cell layer under baseline conditions (*10*), and have different connectivity than superficial neurons (*16*). Granule neurons generated at Gd 4.5 and 7.5 were functionally integrated into the OB circuitry as measured by c-fos expression (Fig. 2G and I). Moreover, in mothers, the increase in new neurons was due to those expressing c-fos (Fig. 2G), predominantly located in the superficial GCL layer (Fig. S2F). Enhanced neuronal activity in analog+ neurons was not caused by different olfactory cues, as the number of c-fos+ analog+ neurons was comparable between virgins alone and virgins exposed to pups for 1 hour (Fig. 2G). Together, this indicates that newborn superficial GCL neurons in mothers are specifically generated in anticipation of and used for maternal care.

In the glomerular layer (GL), specific subtypes namely CalR+ and Calbindin+ (CalB) neurons born at Gd 4.5 and 7.5, and Tyrosine hydroxylase+ (TH) neurons at Gd 7.5, increased during pregnancy (Fig. S4A to E). Newly-generated CalR+ and TH+ neurons were found in clusters extending over 2-to-5 glomeruli (Fig. S4F to H). Unexpectedly, the MCL, a layer into which adult-born neurons rarely integrate in basal conditions, was the only layer in which neurogenesis was increased at Gd 0.5 and Gd 2.5 + 20dpi (around birth) (Fig. 2C and D), as well as at Gd 4.5 and Gd 7.5 +20dpi (early perinatal care) (Fig. 2E and F). These neurons were mature (NeuN+ Doublecortin-), and therefore not migrating through the MCL (Fig. S2G and H). While the increase in interneurons in the MCL born at Gd4.5 and 7.5 in mothers was due to those expressing c-fos, the increase in the Gd0.5/Gd2.5 born cells was mainly due to c-fos-negative neurons (Fig. 2H and I), suggesting that these two neuronal waves may have different functions. Furthermore, neurogenesis was also increased in the AOB (Fig. 2C to F), where adult born-neurons have been implicated in social and reproductive behaviour (*17*). Neurons born at Gd 4.5 and Gd 7.5 were selectively increased in the GCL of the AOB (Fig. S3B and C), and a higher proportion was c-fos-positive in mothers pulsed at Gd 7.5 (Fig. S3D). Thus, pregnancy leads to the recruitment of different pools of NSCs that generate different interneuron subtypes in a temporally coordinated manner, that integrate into the OB at different stages of early motherhood. Depending on their location and their time of addition, the c-fos expression pattern of these neurons differs, highlighting the importance of adult-born olfactory bulb interneuron heterogeneity in specific physiological contexts.

Interestingly, the total number of Olig2+ cells increased in the GCL during the perinatal care period, and decreased around peri-weaning. This transient increase was due to immature NG2+ oligodendrocyte progenitors (OPCs) and not mature oligodendrocytes (Fig. 3A and B), and occurred specifically in the GCL (for EPL and GL see Fig. S5A and B). Birth-dating of Olig2+ cells using a thymidine analog pulse chase revealed that the increase of NG2+ Olig2+ OPCs in the OB slightly preceded the integration of newborn neurons during pregnancy (Fig. 3C-E). We detected no increase of analog+ Olig2+ cells in the OB at early time points after pulsing (Gd4.5+ 3dpi) (Fig. S5C), suggesting the newly generated OPCs were not born locally. Taken together, our data suggest that OPCs born during the first days of pregnancy transiently colonize the GCL, concomitant with the addition of newborn neurons in this layer.

**Fig. 3.**
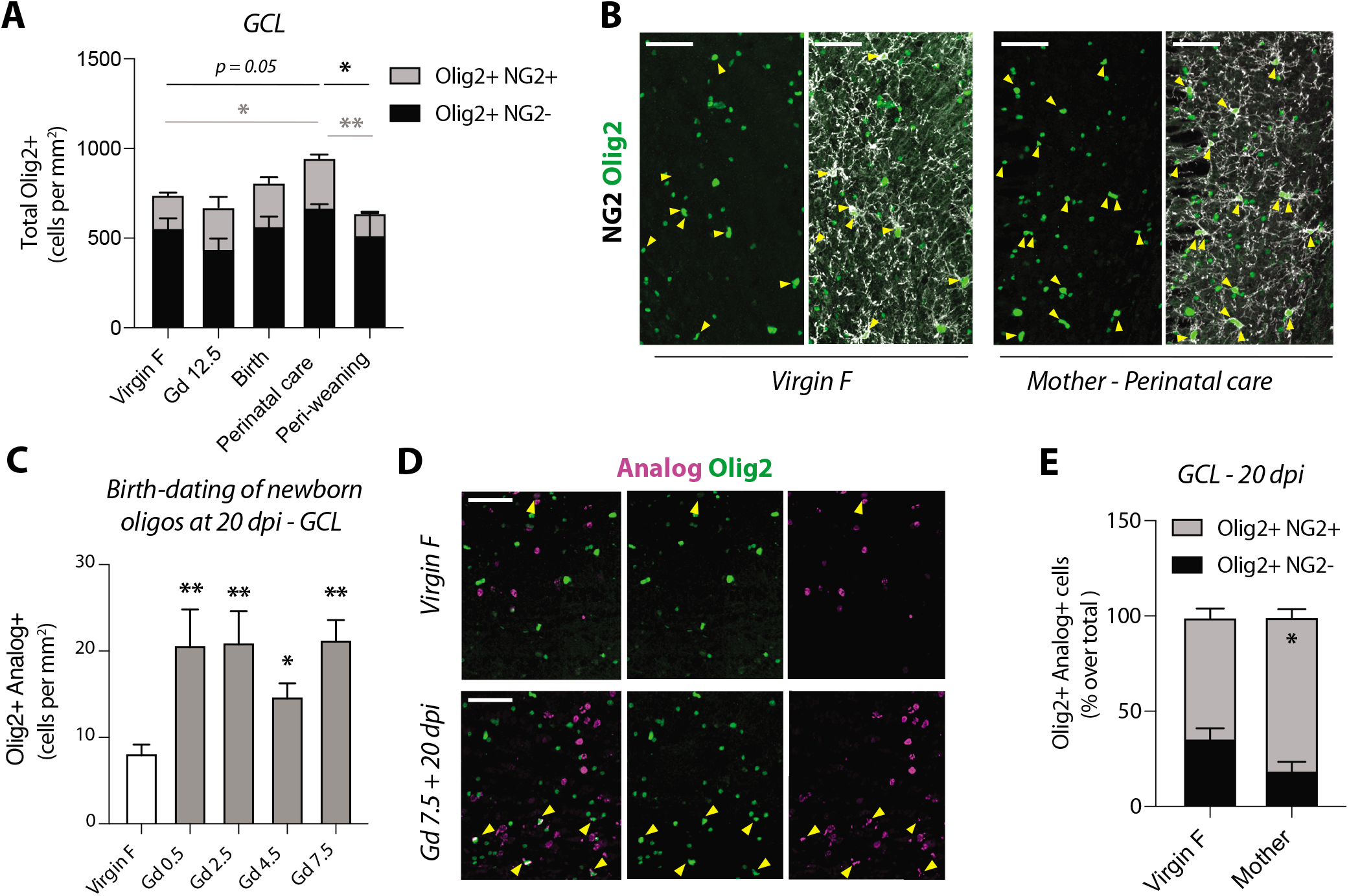
Increase of oligodendrocyte progenitor cells in the GCL during perinatal care period. (**A**) Quantification of total Olig2+ NG2+ oligodendrocyte progenitors (OPCs) and Olig2+ NG2-mature oligodendrocytes in the GCL of the main olfactory bulb. Black lines and asterisks indicate significant differences in total Olig2+ cells (NG2+ and NG2-); grey lines and asterisks indicate differences in Olig2+ NG2+ cells only. (**B**) Representative micrographs to **A** (yellow arrowheads indicate Olig2+ NG2+ cells). (**C**) Quantification of Olig2+ analog+ cells pulsed at different gestation days in the GCL at 20 dpi. (**D**) Micrographs illustrating analog+ Olig2+ cells (yellow arrowheads) in virgin and mothers at perinatal care. (**E**) Proportion of OPCs of total analog+ Olig2+ cells at 20 dpi in virgin female and mothers (Gd4.5 and Gd 7.5 pooled). Scale bars, 50μm.

Adult born-neurons that integrate into OB circuitry persist for many months (*18, 19*). As the neurons generated during pregnancy were specifically increased at key physiological time points during motherhood, we investigated whether they persisted 10 days later, when pups are progressively feeding on solid food and mothers less engaged in maternal care (Fig. 4A). Strikingly, in all layers in which neurogenesis was enhanced in mothers, except the GL, the number of newborn neurons decreased between 20 and 30 dpi (Fig. 4B to D), as did the number of c-fos+ analog+ cells (Fig. S6A to C). This decrease in analog+ cells was not observed in the matched virgin controls (Fig. 4B to D). Interestingly, neurons born at different gestation days and integrated in different layers exhibited different survival rates. Indeed, MCL interneurons born at Gd0.5 had been culled by 30 dpi (Fig. 4B), while some born at Gd7.5 persisted in mothers, beyond weaning (Fig. 4B). Similarly, a small proportion of interneurons in the AOB born at Gd 7.5 also persisted in mothers (Fig. 4D). These may have different functions than those needed in perinatal care, such as potential roles in sexual behavior (*20*). Of note, the dynamics of neuronal addition in the GL were more complex, and the newly generated neurons in the GL were not transient (Fig. S6D-F). Thus, in addition to constitutive neurogenesis leading to long-lasting integration of neurons in the olfactory bulb, substantial targeted and transient neurogenesis occurs during key physiological experiences in the adult brain.

**Fig. 4.**
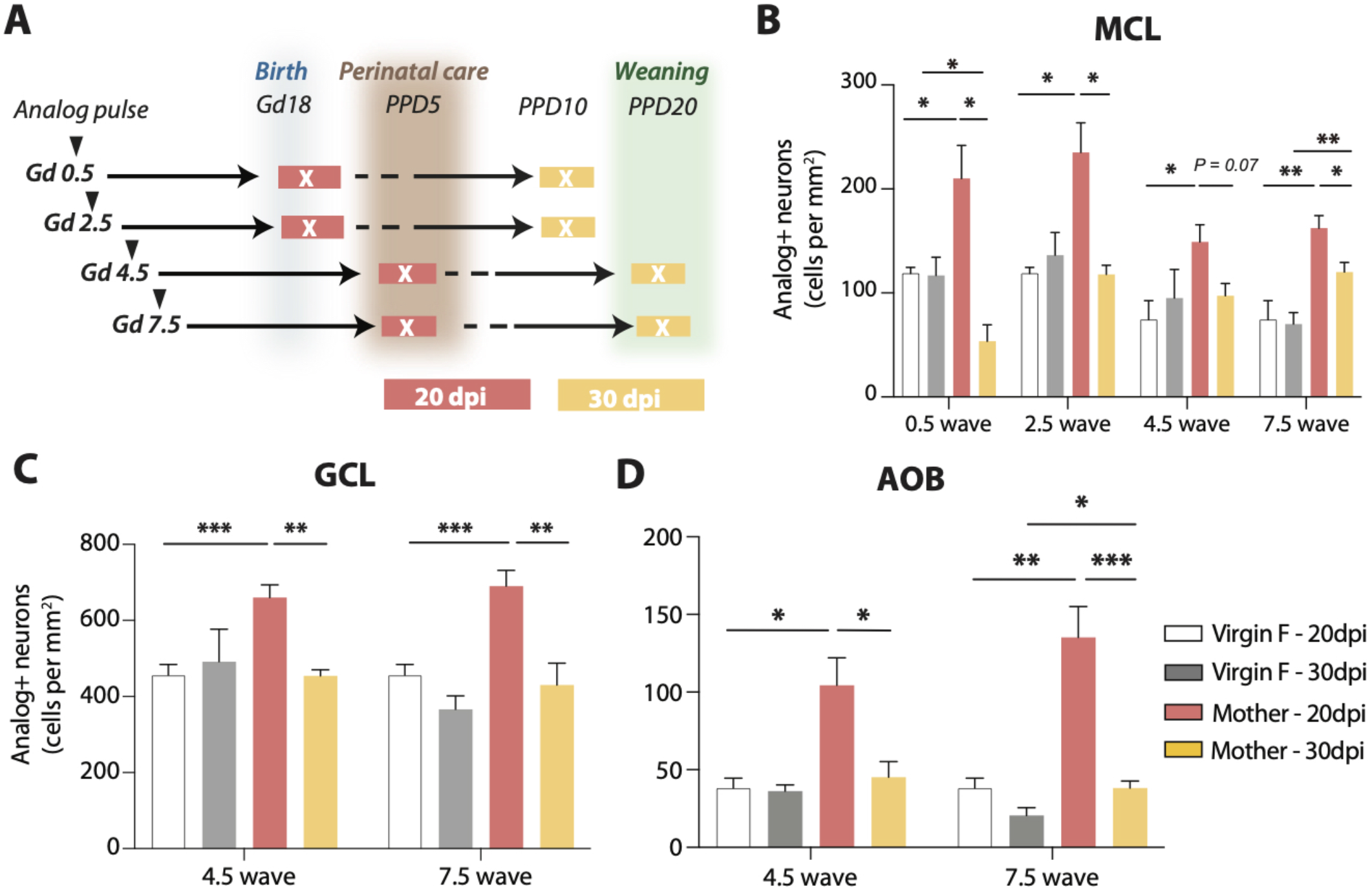
Transient addition of pregnancy-associated interneurons to the olfactory bulb. (**A**) Schema showing relationship of 20 and 30dpi timepoints to birth, perinatal care and weaning for cells born on different gestation days. (**B** to **D**) Dynamics of newly-generated neuronal waves at 20 dpi and 30 dpi in the MCL (**B**) and GCL (**C**) of the MOB and in the AOB (**D**).

Our findings shed new light on NSC heterogeneity in the adult mouse brain, and reveal the existence of stem cell pools that are selectively recruited in different physiological states (Fig. S7). More broadly, this may be linked to the transient generation of distinct neuronal and glial subtypes required for temporally changing physiological needs. In songbirds and chickadees, seasonal neurogenesis has been linked to seasonal song-learning and food-caching (*21, 22*). Here, we identify much shorter time scales over which new neurons and oligodendrocyte progenitors are transiently produced and functionally integrated, which may have important consequences for olfactory bulb plasticity in the adult mammalian brain.

During pregnancy, the choreographed recruitment of regionally-distinct NSC pools at different times is likely due to the complex interplay of local and long-range signals, including innervation and systemic hormonal, as well as metabolic, fluctuations. Importantly, different physiological states, such as hunger and satiety (*3*) and, as shown here, pregnancy, recruit spatially distinct NSC pools, and in turn generate different olfactory bulb interneuron subtypes. The general principle of transient neurogenesis on physiological demand may be conserved across evolution, including in humans, such as during pregnancy where the sense of smell can dramatically change. Altogether, our findings reveal an underlying functional map of adult NSC heterogeneity in the V-SVZ, and suggest that timing, space and physiological context are key to fully decode it.

## Acknowledgments

We thank members of the Doetsch lab, P. Scheiffele and D. Thaler for discussion and comments on the manuscript; and the Biozentrum Imaging Core Facility.

## Funding

This work was supported by Swiss National Science Foundation 31003A_163088 (F.D), European Research Council Advanced Grant (No 789328) (F.D.), the University of Basel, Biozentrum PhD Fellowship University of Basel (C.S.) and the Doris Dietschy und Denise Dietschy-Frick-Stiftung (C.S.).

## Author Contributions

Conceptualization: F.D., Z.C. and C.S.; Performed experiments: Z.C. and C.S.; Data Analysis: Z.C. and C.S.; Supervision: F.D.; Manuscript writing: Z.C., C.S. and F.D.

## Competing Interests

The authors declare no competing interests.

## Notes

### Competing Interest Statement

The authors have declared no competing interest.

