## Supplementary Materials for "Spatio-Temporal Recruitment of Adult Neural Stem Cells for Transient Neurogenesis During Pregnancy"

**This PDF file includes:**

Materials and Methods
Figs. S1 to S7

### Materials and Methods

#### Animal use

Two to three month-old heterozygous *hGFAP::GFP* mice (*1*) were used for timed-matings. Mice of each pregnancy cohort were compared to age-matched virgin controls, injected and processed at the same time. Females from different litters were randomized to form the control virgin and pregnancy/lactation groups. Virgin females were housed in cages of three to four. To set up the timed mating, two females were placed in a cage together with one male 30min before onset of the dark cycle. On the morning of the copulatory plug, mated females were transferred into fresh cages and housed in pairs until the end of experiment. The females spent no more than 3 days with a male before separation, which is important in our context, as male pheromones have been described to have an effect on V-SVZ neurogenesis after seven but not two days of exposure (*2*). Mice kept for the 30dpi timepoints were housed alone with their litter shortly after they had given birth. The morning of the plug is defined as Gd 0.5. This mouse strain gives birth between gestation days 18.5-19.5.

For experiments assessing c-fos in the OB, three groups were used: virgins with no contact to pups, virgins exposed to pups for 1 hour, and mothers with their pups. All groups (virgins and mothers with pups) were transferred to fresh cages and allowed to acclimate for 1 hour. For the virgins exposed to pups for 1 hour group, pups were added to the cage for 1 hour. All mice were sacrificed after 1 hour.

Mice were housed in a 12:12h dark/light cycle with ad libitum access to food. All experimental cohorts were sacrificed at the same time of the day, i.e the morning. All experiments were performed under a license approved by the veterinary office of canton Basel-Stadt.

#### Thymidine analog injection

Virgin control females, pregnant and lactating mice were pulsed with a single intraperitoneal injection of thymidine analog on specific days of gestation and mice sacrificed 20 or 30 days later. All injections were performed in the morning. The following timepoints were investigated: Gd 0.5 (morning of copulatory plug), Gd 2.5, Gd 4.5 (day of blastocyst implantation), Gd 7.5. Three different thymidine analogs were used at equivalent stochiometric doses: 1) 5-bromo-2`deoxyuridine (BrdU; Sigma B5002-5G; 50mg/kg body weight), 2) 5-Chloro-2`deoxyuridine (CldU; Sigma C6981-100MG; 43mg/kg body weight), and 3) 5-Iodo-2`deoxyuridine (IdU; Sigma I7125-5G; 58mg/kg body weight). For all three thymidine analogs, 10mg/ml stock solutions were prepared in sterile 0.9%NaCl. A few drops of 5M NaOH were added to the IdU solution to dissolve it. Aliquots of each solution were stored at -20°C (BrdU, CldU) or refrigerated (IdU) up to 6 months. Solutions were vortexed at 37°C for 10-15min prior to injection.

All pregnant mice were compared to matched controls injected with the same thymidine analogs.

Gd 0.5/Gd 2.5 + 20 dpi cohorts and their matched virgin control mice were co-injected with CldU at Gd0.5 and IdU at Gd 2.5. Similarly, Gd 4.5/Gd 7.5 + 20 dpi cohorts and their matched virgin control mice were co-injected with CldU at Gd4.5 and IdU at Gd 7.5. For the 30dpi time point, Gd 2.5/Gd 4.5 cohorts and their matched virgin control mice were co-injected with CldU at Gd2.5 and IdU at Gd 4.5. Gd 0.5+ 30 dpi (CldU) and Gd7.5 + 30 dpi (IdU or BrdU) cohorts were single injected. Mice were sacrificed 20 or 30 days after the last injection of analog.

**Tissue preparation**

Mice were deeply anaesthetized with pentobarbital and perfused with 0.9% saline followed by ice-cold 3.2% paraformaldehyde (PFA, Electron Microscopy Sciences) in 0.1M phosphate buffer (PB). Brains were postfixed in 3.2% PFA for 24h at 4°C, washed 3x in PBS over a day and subsequently stored at 4°C in 0.05% PBS-azide until processing. Coronal sections were cut on a vibrating microtome (Leica VT10006). Olfactory bulbs were sectioned at 30µm (OB) and the rest of the brain at 25µm.

V-SVZ sections were serially collected in sequential wells of a twelve-well plate and stored in 0.05% PBS-azide until processing. Olfactory bulb sections were cut at 30µm thickness between Bregma +5.2 mm to +3.0 mm and were serially collected in different wells.

**Immunostaining**

Immunostaining of virgin and pregnant /lactating groups was performed on bregma-matched sections. Floating sections were blocked for 1h at RT in PBS with 0.05% (for $\alpha$EGFR, $\alpha$NG2) or 0.3-0.5% (for all other antibodies) Triton-X 100, 10% normal donkey serum (Gene Tex, #GTX73245). The sections were then incubated in the primary antibody solution prepared in the same blocking solution at 4°C overnight, washed 3x15min in 1X PBS before incubating for 1-2h at RT in the secondary antibody prepared in blocking solution. Finally, the sections were washed 3x15min, and counterstained with DAPI. Sections were mounted on glass slides using Aqua-Poly/Mount (Polysciences #18606-20).

When combining thymidine analog detection with other immunostaining, other antibodies were immunostained first, sections fixed for 10 min at RT in 3.2% PFA, and washed 3x15min in PBS. Sections were then incubated for 20 to 30 min in 2M HCl at 37°C. After removing the HCL, the pH was neutralized using a 0.1M Borax pH 8.5 solution for 10min at RT, followed by 3x15min wash in 1X PBS. Detection of the thymidine analogs was performed with the following antibodies: Anti-BrdU rat (abcam) for detecting BrdU and CldU, and anti-BrdU mouse (clone B44, BD-Bioscience) for IdU. In mice co-injected with both CldU and IdU, we distinguished the two analog types following the protocol described in Podgorny et al. 2018 (*3*), except we shortened the DNA denaturation step to 20-30 min. Single injected controls were processed in parallel to set the imaging settings and thresholds.

**Antibodies**

The following primary antibodies were used: anti-BrdU (rat, 1:500-1:1000, abcam #ab6326); anti-BrdU (mouse clone B44, BD-Bioscience), anti-Calbindin (rabbit, 1:1000, Chemicon, #AB1778), anti-Calretinin (rabbit, 1:1000, SWANT, #7697), anti-Calretinin goat, 1:1000, Labome Millipore, #ab1550), anti-Tyrosine Hydroxylase (sheep, 1:500, Millipore, #AB1542), anti-doublecortin, DCX (goat, 1:100, Santa Cruz, discontinued); anti-doublecortin, DCX (guinea pig, 1:1000, MerckMillipore, #ab2253), anti-EGFR (goat,1:100, R&D, #BAF1280); anti-EGFR (rabbit,1:100, abcam, # ab52894), anti-GFAP (chicken, 1:600, Millipore, #MAB5541), anti GFAP (rat, 1:600, Invitrogen, #13-0300), anti-Ki67 (rabbit, 1:100, abcam, #ab15580), anti-NeuN (mouse, 1:100, Millipore, #MAB377); anti-NeuN (rabbit, abcam, #ab177487); anti-NG2 (rabbit, 1:100, Millipore); anti-Olig2 (rabbit, 1:150, Millipore, #ab9610); anti-Olig2 (goat, 1:150, R&D, #AF2418), anti-c-fos (rabbit, 1:1000, Synaptic Systems, 226003).

The following secondary antibodies were used: Alexa Fluor-conjugated (405, 488 and 647; 1:200-1:600, Invitrogen) and Cy3-conjugated (1:600-1:1000, Jackson ImmunoResearch) Fab2 fragments raised in donkey.

**Image acquisition and quantification**

Images were acquired using LSM700, LSM800 and LSM880 confocal microscopes (Zeiss), the confocal spinning disk microscope SpinSR (Olympus) or the Zeiss AxioScan Z1. On the LSM700 and LSM800, images were acquired with PLAN APO 25x/0.80NA and PLAN APO 40x/1.3NA objectives. On the LSM880, a PLAN APO 40x/1.2 NA objective was used, and on the SpinSR a UPL S APO 30x/1.05NA. Tile scans of the entire OB and V-SVZ were acquired, with z-stacks encompassing the entire thickness of the sections. Individual z step sizes were 1-2µm, depending on the staining. Images of the same immunostaining were acquired on the same microscope.

Analysis of proliferating NSCs (Fig.1B and C) was performed on sections between Bregma +0.3 mm and 0 mm. For each bregma, two to four sections per mouse were analyzed for n=3-6 mice. Cells were quantified manually using the counter tool in Fiji. Data are presented as densities (cells per mm3 for NSCs in the V-SVZ, or cells per mm2 for MOB and AOB sections).

Counts in the OB were performed on entire olfactory bulb sections. For the quantification of cells in the main olfactory bulb GCL and GL, two sections were used per mouse for n=3-7 mice per group. For the MCL and AOB, four to five sections per animal were sampled due the low number of total analog cells incorporated in these regions. For quantification of NeuN+ analog+ cells in the superficial and deep GCL only (Fig. S4A), two separate strips on the medial and lateral olfactory bulbs capturing the entire dorsal-ventral length were quantified.

**Statistical analysis**

Significance was established using two-tailed, non-parametric student's *t*-tests for pair-wise comparisons. 2-way ANOVA was used for testing the pregnancy effect, and time effect for all quantification of NSC dynamics. Significance was established at *p<0.05, **p<0.01, ***p<0.001. Error bars indicate the standard error of the mean (SEM). Statistics were performed using Prism 6 software.

For figures 2**C** to **F** (fold change quantifications), significance was calculated on raw values relative to the matched virgin control group. In figures 2**G** and **H** (c-fos quantifications), statistical tests were comparing "plain mothers with pups" to the pooled group of virgins including both "virgin females exposed to pups" and "virgin females without pup exposure".

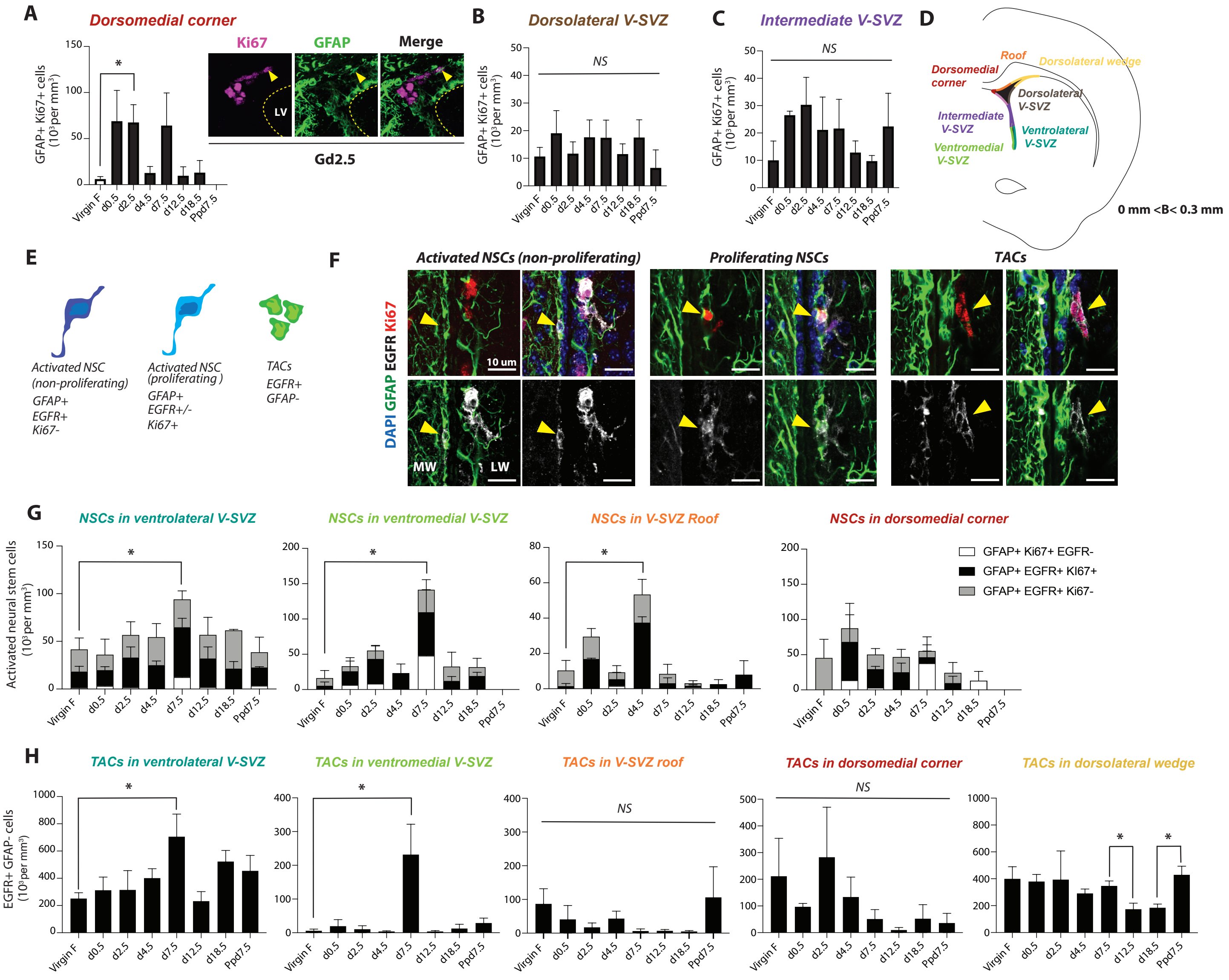


**Fig. S1. Regional dynamics of adult NSCs and their progeny in the V-SVZ during pregnancy (A-C)** Quantification of GFAP+ Ki67+ cells in the dorsomedial corner and corresponding GFAP/Ki67 images at Gd 2.5 (**A**), in the dorsolateral V-SVZ (**B**), and intermediate V-SVZ (**C**). (**D**) Schema illustrating different V-SVZ domains analyzed. (**E**) Schema of markers used to quantify activated NSCs and transit-amplifying cells (TACs). (**F**) Representative micrographs of non-proliferating activated NSCs, proliferating activated NSCs, and transit amplifying cells (EGFR+ GFAP-). (**G**) Quantification of activated and dividing NSCs in different V-SVZ domains (GFAP+ EGFR+ Ki67-, GFAP+ EGFR+ Ki67+, GFAP+ Ki67+ EGFR-). Asterisks indicate significant differences in total cell numbers represented in the stacked bar graphs. (**H**) Quantification of transit-amplifying cells in different V-SVZ domains.

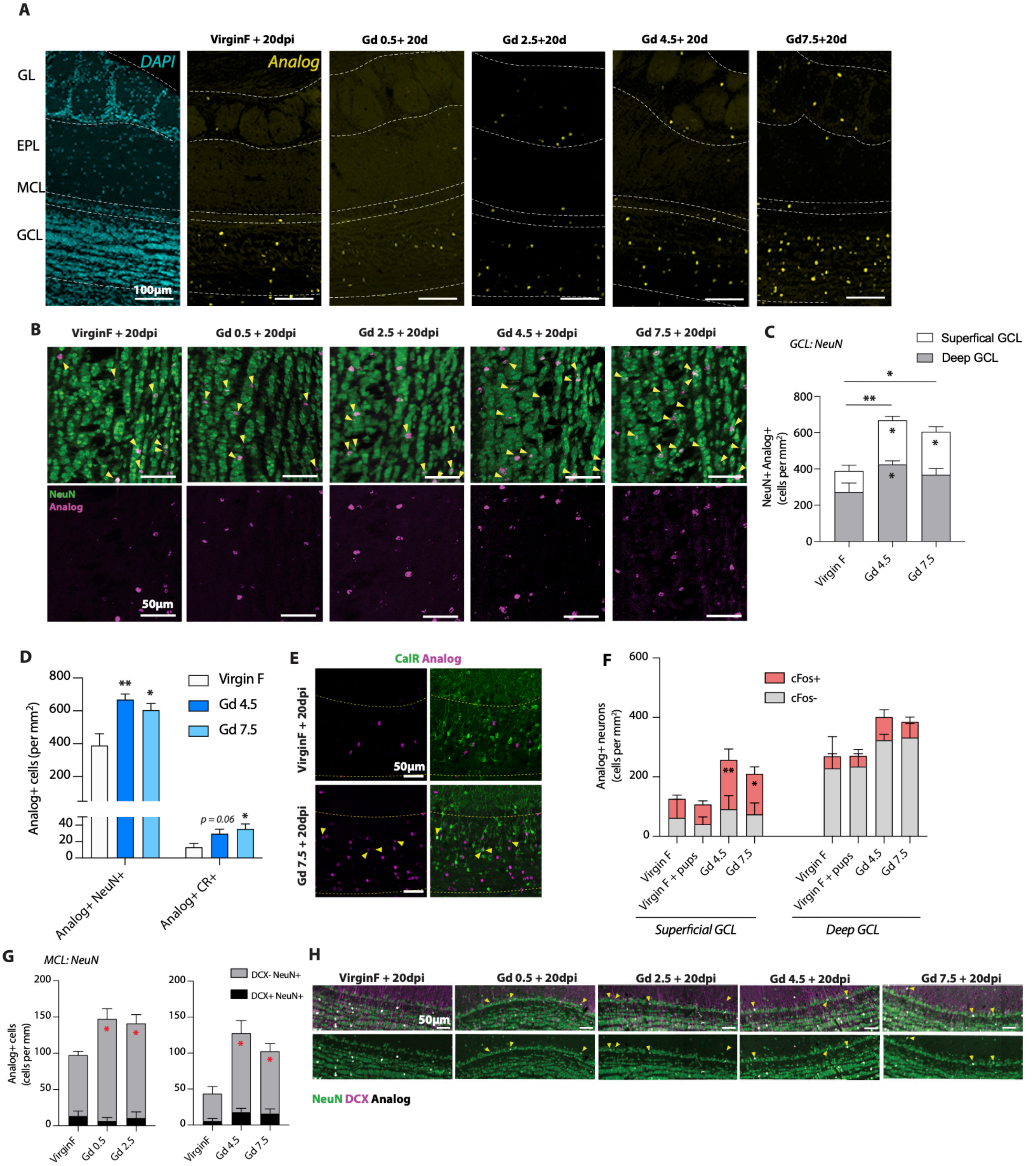


**Fig. S2. Characterization of newborn neurons in the GCL and MCL of the MOB at 20 dpi.** (**A**) Left panel: Low power image of DAPI stained main olfactory bulb showing different layers. Right panels: Images of thymidine analog+ cells in the MOB of virgin mice and mothers pulsed at Gd 0.5, Gd 2.5, Gd 4.5 or Gd 7.5. (**B** to **E**), Characterization of newborn cells in the GCL. (**B**) Representative images of GCL showing NeuN in green and analog in magenta. (**C**) Quantification of newborn NeuN+ neurons in both superficial and deep GCL at Gd 4.5 and Gd 7.5 + 20 dpi. (**D**) Quantification of total NeuN+ analog+ neurons and CalR+ analog+ neurons in the GCL at Gd 4.5 and Gd 7.5 + 20 dpi. (**E**) Representative images of (**D**), showing CalR in green and analog in magenta. (**F**) Quantification of c-fos+ analog+ neurons in superficial and deep GCL at Gd 4.5/7.5 + 20. Stars inside pink bar graphs show significant differences in c-fos+ analog+ cell number between virgins and mothers. (**G** and **H**) Characterization of newborn cells in the MCL. (**G**) For all pulsed timepoints, the vast majority of analog+ cells in the MCL were NeuN+ DCX-. (**H**) Representative pictures of (**G**), showing NeuN in green, analog in white and DCX in magenta. GL: Glomerular layer; EPL: External plexiform layer, MCL: Mitral cell layer, GCL: Granule cell layer.

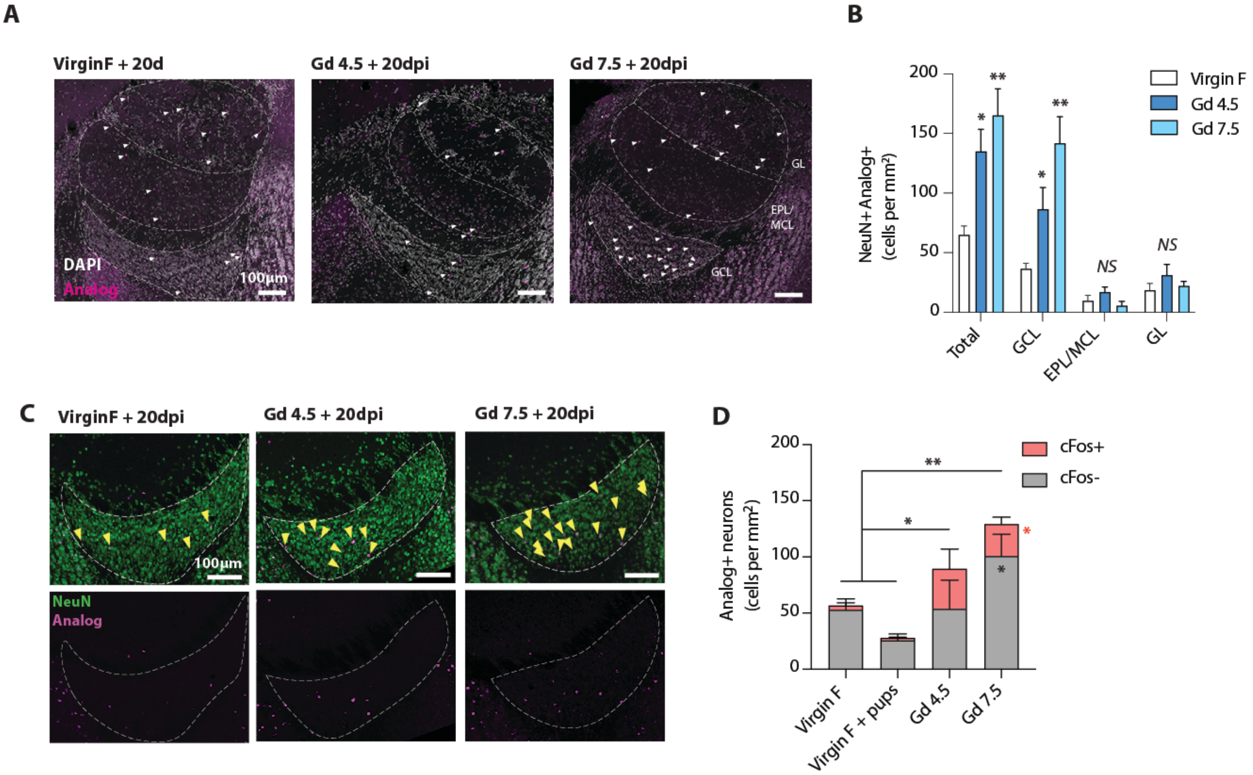


**Fig. S3. Characterization of newborn neurons in the AOB at 20 dpi.** (**A**) Low power images showing analog staining (magenta) in all AOB layers, and DAPI in white. (**B**) Quantification of newborn neurons (NeuN+ analog+) generated at Gd 4.5 and 7.5 in each AOB layer. (**C**) Representative pictures to (B), showing NeuN in green and analog in magenta. Arrowheads indicate double-labeled cells. (**D**) Proportion of c-fos+ analog+ neurons in the AOB at Gd 4.5/7.5 + 20 dpi. Red star shows significant increase in c-fos+ analog+ cell number in Gd7.5 mothers compared to pooled virgins. Star inside the grey bar graph shows significant increase in c-fos- analog+ cells in Gd7.5 mothers compared to pooled virgins. GL: Glomerular layer; EPL: External plexiform layer, MCL: Mitral cell layer, GCL: Granule cell layer.

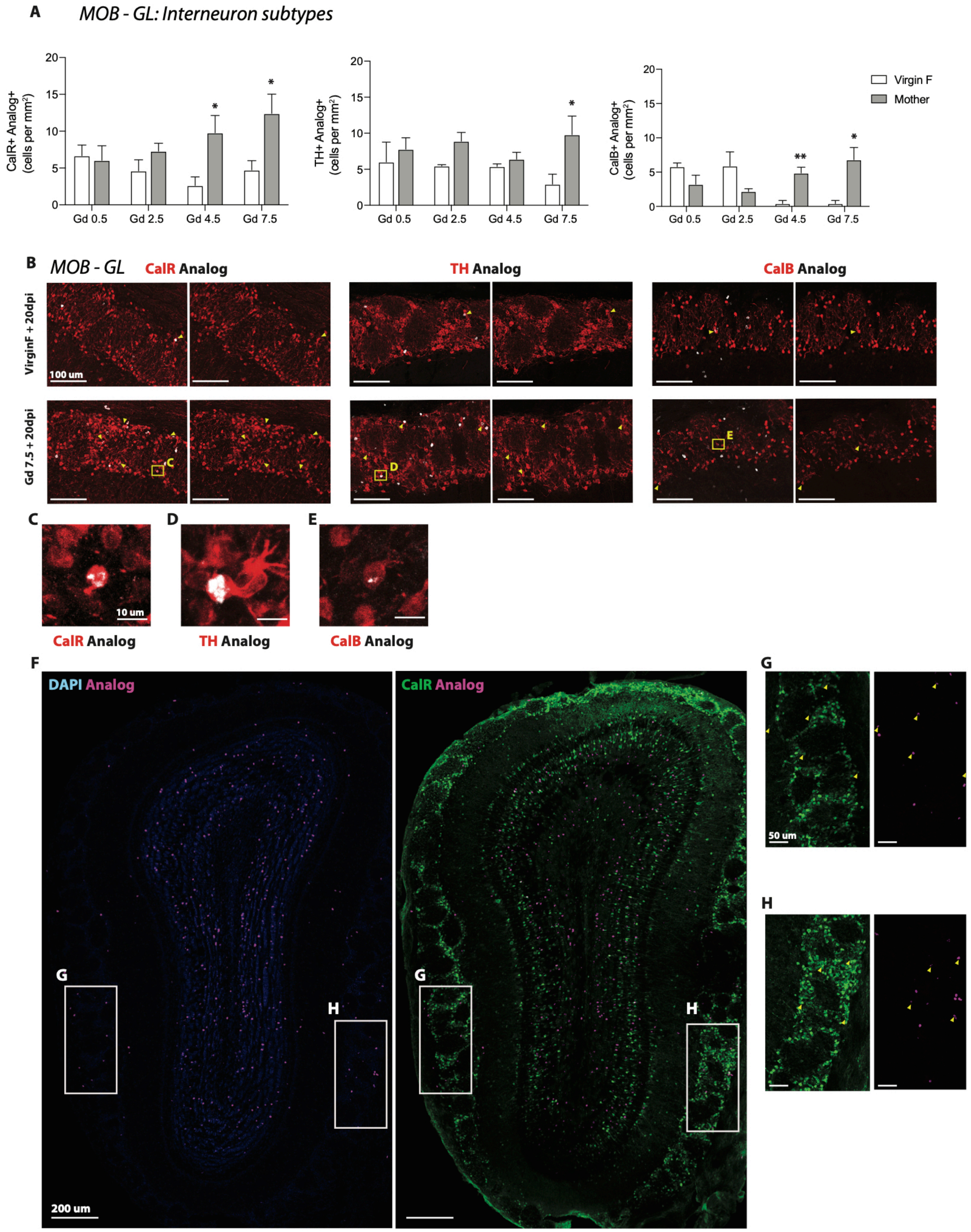


**Fig. S4. Characterization of newborn neurons in the glomerular layer of the MOB at 20dpi.** (A) Quantification of analog+ CalR+, TH+ and CalB+ cells compared to their corresponding matched virgin controls. (**B**) Representative images of analog+ CalR+, TH+ and CalB+ subtypes in the GL comparing virgin females (top panels) and Gd 7.5 + 20 dpi (bottom panels). (**C** to **E**), High-power images of CalR+, TH+ and CalB+ analog double-positive cells shown in boxes in (**B**). (**F**) Low power view of an entire OB coronal section in a Gd7.5 + 20 dpi mother, showing clusters of glomeruli containing CalR+ analog+ cells in boxes. (**G** and **H**) are high-magnification images corresponding to boxed GL regions in (**F**).

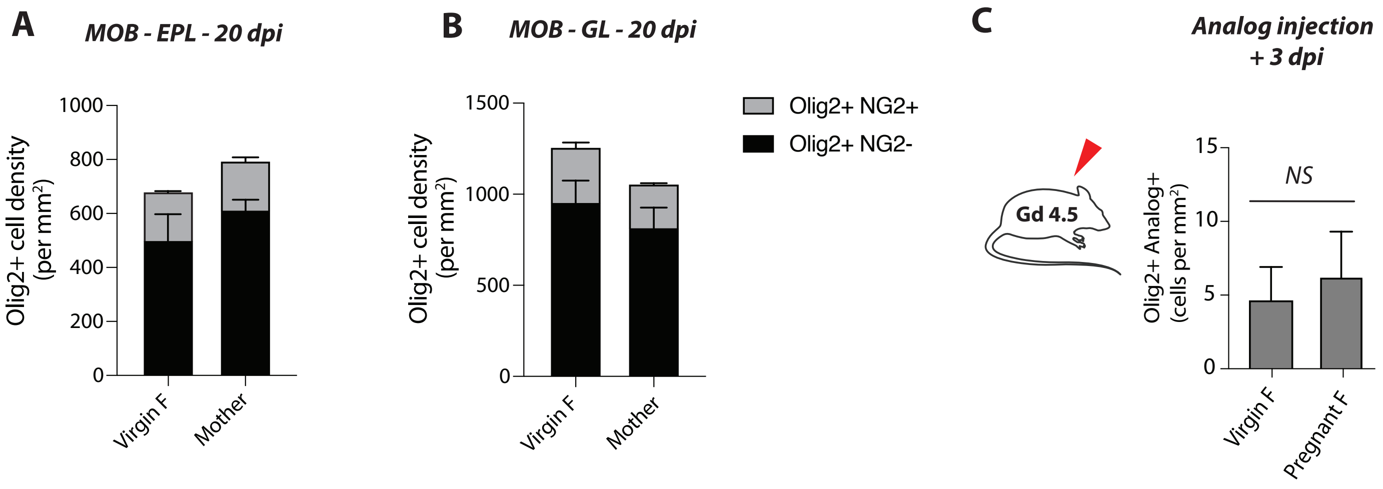


**Fig. S5. Increased generation of OPCs during perinatal care period is specific to the GCL.** (**A** and **B**) Quantification of total Olig2+ NG2+ (oligodendrocyte progenitor, OPCs) and Olig2+ NG2- (mature oligodendrocyte) cells in the EPL (A), and in the GL of the MOB (**B**). (**C**) Quantification of analog+ Olig2+ cells in the GCL in pregnant females pulsed at Gd 4.5, and analyzed 3 days later, at Gd 7.5. MOB: Main olfactory bulb, GL: Glomerular layer, EPL: External plexiform layer.

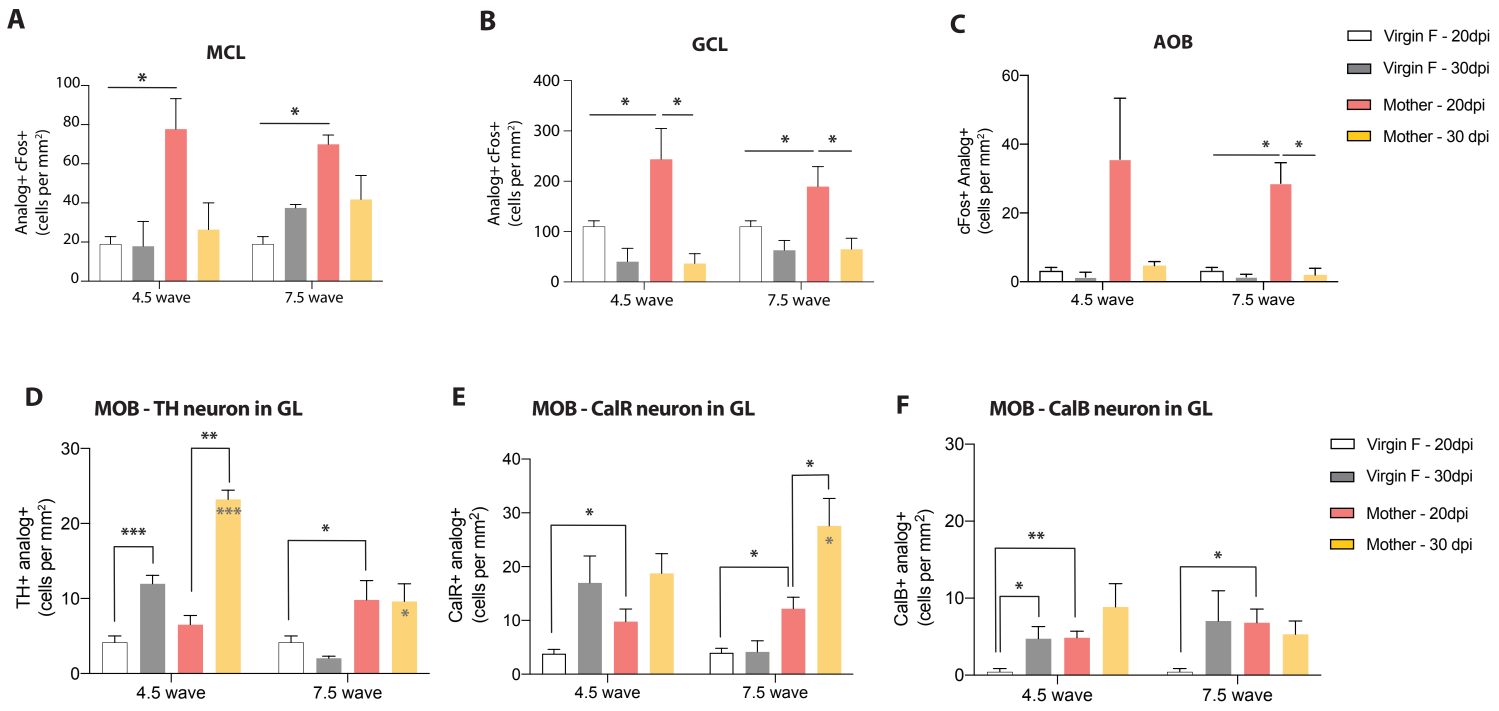


**Fig. S6. Dynamics and activity of newborn neurons in the olfactory bulb.** (**A** to **C**) Quantification of c-fos+ analog+ cells in the MCL of the MOB (**A**), the GCL of the MOB (**B**), and the AOB (**C**) at 20 *vs* 30 dpi. (**D** to **F**). Quantification of dynamics of TH+, CalR+ and CalB+ newborn neurons in the GL at Gd 4.5/Gd 7.5 + 20dpi and 30dpi. Grey stars inside the yellow bars show significant differences as compared to the virgin 30dpi.

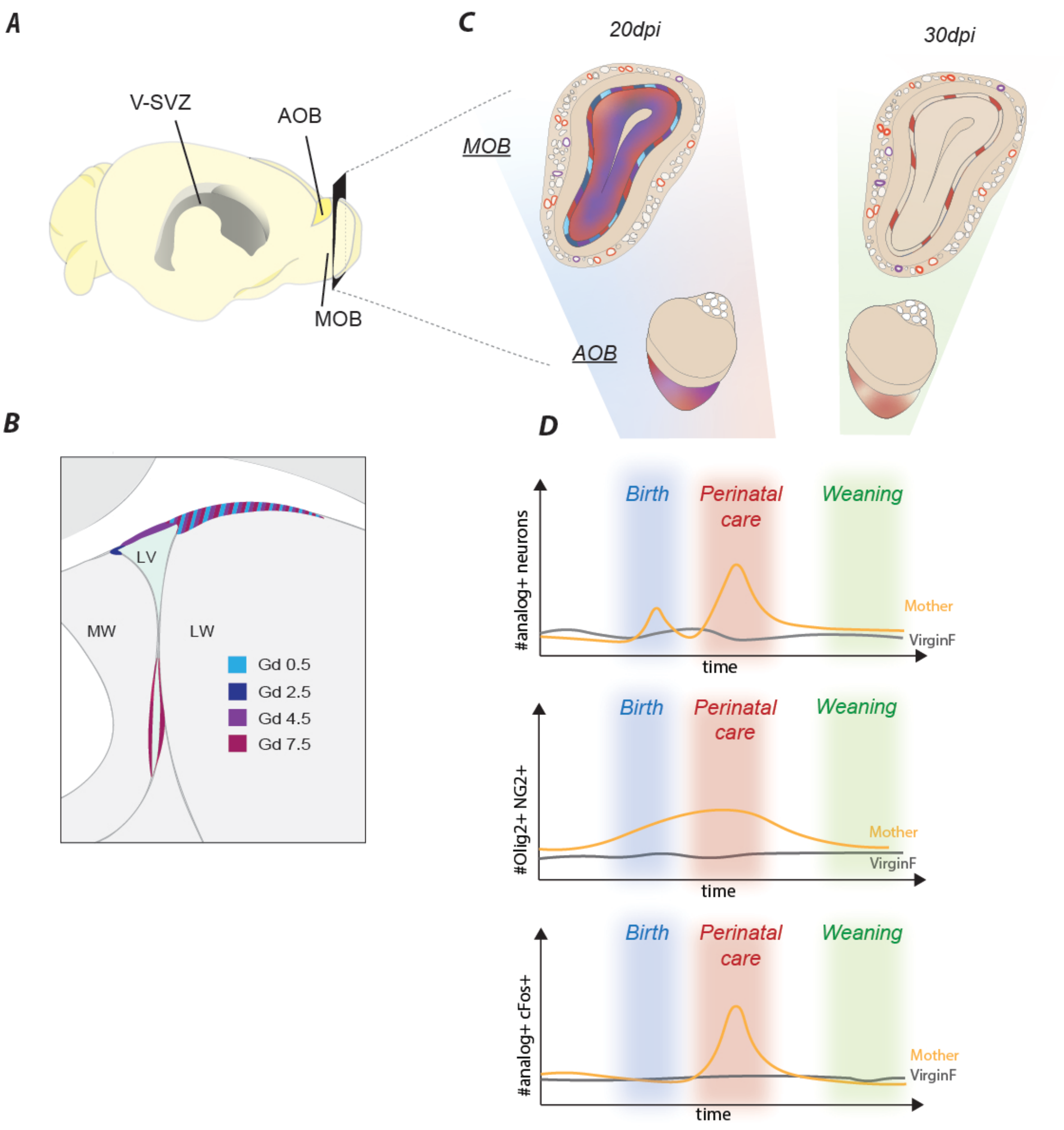


**Fig. S7. Spatiotemporal stem cell recruitment during pregnancy results in transient addition of newborn neurons at birth and during perinatal care period.**

(**A**) Schema of mouse brain showing location of V-SVZ adjacent to the lateral ventricles (grey). (**B**) Summary schema of V-SVZ domains recruited at different days of gestation. (**C**) Schemas of main and accessory olfactory bulbs showing layers with increased neuronal addition at 20dpi of cells labeled at Gd0.5 (turquoise), Gd2.5 (blue), Gd4.5 (purple) and Gd7.5 (red). The majority of these cells are transiently integrated in the OB. (**D**) Schemas summarizing the temporal dynamics of pregnancy-associated interneurons and OPCs in the OB, and transient increase in c-fos+analog+ neurons. LV, lateral ventricles.
